# SRSF3 confers selective processing of miR-17-92 cluster to promote tumorigenic properties in colorectal cancer

**DOI:** 10.1101/667295

**Authors:** Madara Ratnadiwakara, Rebekah Engel, Thierry Jarde, Paul J McMurrick, Helen E Abud, Minna-Liisa Änkö

## Abstract

Almost a half of microRNAs (miRNAs) in mammalian cells are generated from polycistronic primary transcripts encoding more than one miRNA. Mature miRNAs from polycistronic clusters frequently regulate complementary sets of target mRNAs. How the processing of individual miRNAs within the clusters is controlled to give rise to distinct miRNA levels *in vivo* is not fully understood. Our investigation of SRSF3 (Serine-Arginine Rich Splicing Factor3) regulated noncoding RNAs in pluripotent cells identified miR-17-92 cluster as a key SRSF3 target, SRSF3 binding to the CNNC motif 17-18nt downstream of the miRNA stem loop. Here we show that SRSF3 binding site context, not merely the distance from the stem loop, within primary transcript is a critical determinant of the processing efficiency of distinct miRNAs derived from the miR-17-92 cluster. SRSF3 specifically enhanced the processing of two paralog miRNAs, miR-17 and miR-20a, targeting overlapping mRNAs including the cell cycle inhibitor CDKN1A/p21. Functional analysis demonstrated that SRSF3 inhibits CDKN1A expression and promotes cell cycle and self-renewal through the miRNA processing pathway both in normal pluripotent stem cells and cancer cells. Strikingly, analysis of colorectal cancer tumour-normal pairs demonstrated that the SRSF3-regulated miRNA processing pathway is present in a large proportion of colorectal cancer patients and distinguishes poorly differentiated high-grade tumours. Our research uncovers a critical role of SRSF3 in selective processing of miR-17-92 miRNAs, which mechanistically and functionally links SRSF3 to hallmark features of cancer.

## Introduction

MicroRNAs (miRNA) are small non-coding RNAs ~19-25nt in length that play a critical role in post-transcriptional gene regulation. By base-pairing with complementary sequences in their target mRNAs, miRNAs cause translational repression or mRNA cleavage leading to reduced mRNA and/or protein levels (1). Genetic, biochemical and computational studies have identified hundreds of miRNAs in humans with diverse and often conserved functions affecting most developmental process (1, 2). Accordingly, mice with defects in the miRNA biogenesis pathway die during embryonic development (3–5). Loss-of-function of individual miRNAs leads to diverse phenotypes in various organ systems, reflecting the limited expression of many miRNAs during specific stages of development or in distinct cell types (1). Dysregulated miRNA expression is associated with many human diseases, with a strong link between altered miRNA levels and cancer (6). MicroRNAs can affect many cancer hallmarks including sustained cell proliferation, resistance to apoptosis and capacity to metastasise (7).

MicroRNA biogenesis is a tightly regulated multi-step process (8). Post-transcriptional steps involve two consecutive cleavages of the miRNA stem-loop by the RNAseIII enzymes Drosha in the nucleus and Dicer in the cytoplasm (1). The nuclear microprocessor complex consisting of Drosha and two molecules of its co-factor DGCR8 cleaves the pri-primary (pri-)miRNAs to give rise to a ~60nt stem-loop pre-miRNAs (9, 10). Specific sequence motifs within the pri-miRNA have been shown to affect the pri-miRNA cleavage efficiency, including the apical GUG/UGU, a GC and UG motifs 13nt and 14nt upstream of the Drosha cleavage site, respectively, and a CNNC motif 16-18nt/16-24nt downstream of the Drosha cleavage site (11–15). The CNNC motif was previously identified as the consensus binding motif for Serine-arginine rich splicing factor 3 (SRSF3), an RNA binding protein (RBP) with multiple functions in mRNA metabolism (16, 17). In subsequent studies, binding of SRSF3 to the CNNC motif 17-18nt downstream of the Drosha cleavage site was shown to enhance the processing of primiR-30a/c and miR-16 (13, 15). As 60% of the human miRNAs conserved to mouse contain a CNNC motif within this window (13), SRSF3 may play a broad role in modulating efficiency of pri-miRNA processing.

MicroRNAs are located both between and within genes as single miRNA genes or as miRNA clusters producing polycistronic pri-miRNA transcripts with multiple miRNA stem-loops. Polycistronic miRNAs account for ~50% of all miRNAs in mammals (18) and often contain paralog miRNAs of the same miRNA family targeting similar sets of mRNAs (19). The target mRNAs of a miRNA family tend to encode proteins in a common pathway, allowing efficient gate-keeping of cellular processes through miRNA regulation (20). Several miRNA clusters have been found to be essential for normal development and play a role in disease pathology (19). Although polycistronic miRNAs are critical in human development, homeostasis and disease, we know surprisingly little of how their processing is regulated post-transcriptionally. Importantly, miRNAs produced from a common primary transcript are present at highly varying levels in cells and tissues, suggesting that the processing of individual miRNAs within the cluster can be differentially regulated (11, 21, 22). For example, hnRNPA1 was shown to specifically bind to the conserved terminal loop of miR-18a within miR-17-92 cluster and promote its processing independent of other miRNAs of the cluster (22).

The mature miRNA sequences and their organization of miR-17-92 cluster located on chromosome 14 in mouse and on chromosome 13 in human are fully conserved in all vertebrates (23). The primary transcript includes six mature miRNAs: miR-17, miR-18a, miR-19a, miR-19b-1, miR-20a, and miR-92a-1. Dysregulation of miR-17-92 miRNAs has been reported in various cancers, including colorectal cancer (CRC) (23–25), and studies in mouse models and human cell lines have identified the cluster as an oncogene, hence it is also called OncomiR-1 (26). The miRNAs produced from miR-17-92 and its paralog clusters miR-106a-363 and miR-106b-25 are classified into four miRNA families based on their seed sequences: the miR-17, miR-18, miR-19 and miR-92 families (see Fig. 1*D*). Genetic studies in mice have shown that the loss of miR-17-92 leads to perinatal death, while the ablation of miR-106a-363 and/or miR-106b-25 results in no obvious phenotype (27). This suggests that while there is a functional cooperation between the three clusters, miR-17-92 may control pathways that are critical for cell survival. This has also implications to the tumour promoting role of miR-17-92 which mainly involves promoting cell proliferation and cell cycle progression (28, 29). The regulation of cell cycle by miR-17-92 is also fundamental for embryonic stem (ES) cell self-renewal and proliferation (30), suggesting a broad significance in self-renewing cells.

**Fig 1.**
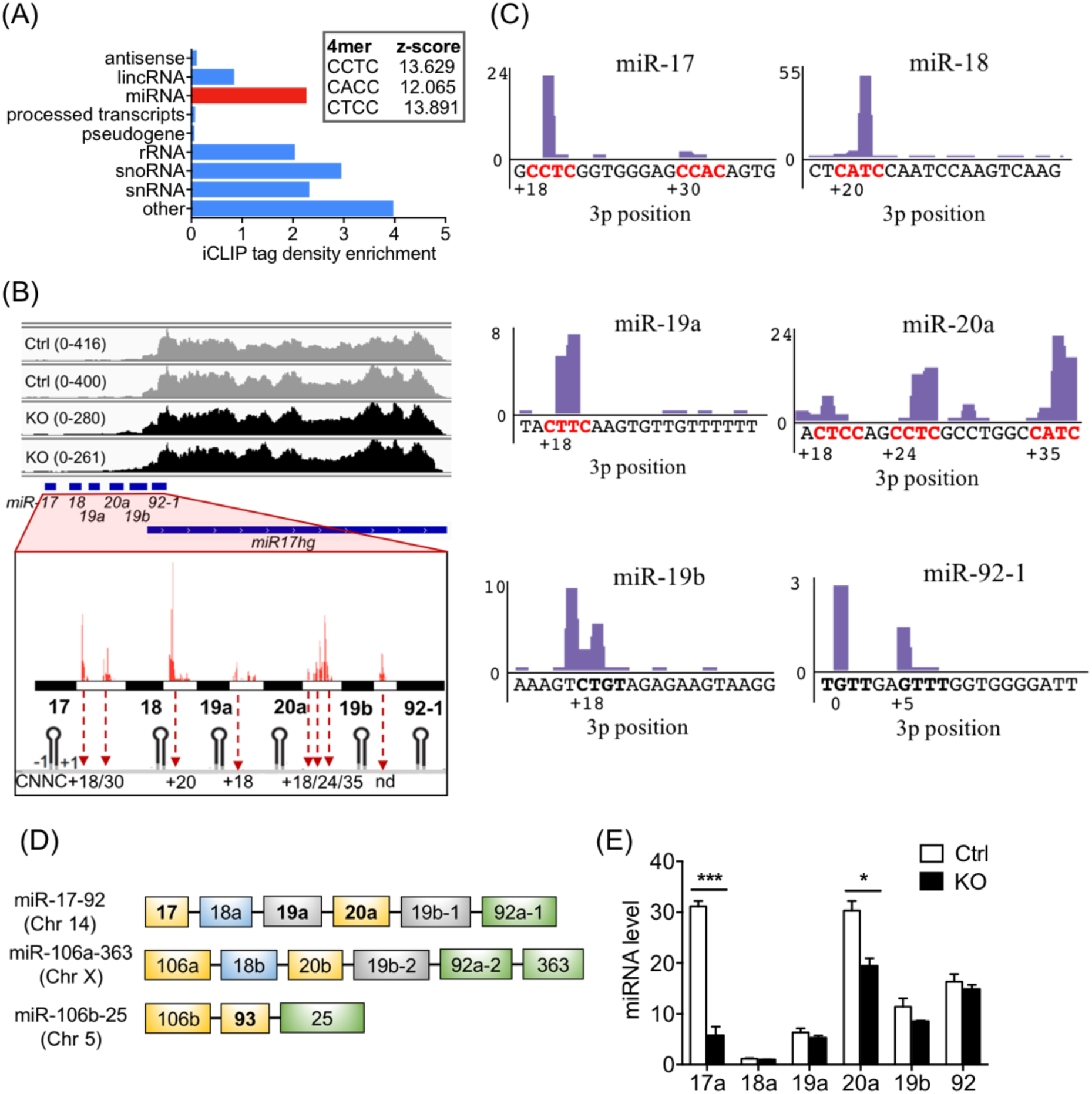
SRSF3 regulates differential processing of miR-17-92 miRNAs in pluripotent stem cells. *(A)* Distribution of significant SRSF3 crosslink sites (FDR < 0.05) over noncoding transcript types. *Inset:* top three enriched tetramers around SRSF3 crosslink sites within miRNA transcripts. *(B)* Expression of *miR17hg* transcript in *Srsf3*-KO and control iPSCs. Window heights are scaled to maximum read counts. *Inset:* An overview of SRSF3 iCLIP binding peaks within pri-miR-17-92 cluster. CNNC motifs within SRSF3 binding peaks are marked with red arrows. *(C)* SRSF3 iCLIP binding peaks within the individual miRNAs of miR-17-92 cluster in mouse ES cells. The 3p position is counted from the 3’ Drosha cleavage site as described in (13). The CNNC motifs are marked in red colour. *(D)* Schematic illustration of three paralog clusters miR-17-92, miR-106a-363, and miR-106b-25. The colours mark miRNAs of the same family. MicroRNAs bound by SRSF3 at 16-18 nt downstream of the Drosha cleavage site are in bold. *(E)* TaqMan analysis of the mature miRNAs of miR-17-92 cluster in *Srsf3*-KO and control iPSCs (*p<0.05, ***p<0.0005, Unpaired Student’s t-test, two-tailed, data as mean ± SEM, n=3). See also Fig. S1

We have previously demonstrated that SRSF3 is essential for self-renewal in mouse pluripotent stem cells (17). SRSF3 enhanced cell proliferation during somatic reprogramming and in pluripotent cells similar to that observed in various human cancer cell lines (17, 31, 32). However, the direct molecular mechanisms underlying the cell proliferation promoting function of SRSF3 have remained elusive although some candidate RNAs have been proposed (31, 32). Here we demonstrate that SRSF3 binding to the CNNC motif of miR-17 and miR-20a, two paralog miRNAs derived from the miR-17-92 polycistron, leads to their enhanced processing and altered expression of the miRNA target mRNAs encoding key cell cycle regulators in mouse pluripotent cells, human cancer cell lines and primary colorectal tumours. We demonstrate how SRSF3 mediated selective processing of miR-17 and miR-20a enhances self-renewal potential, distinguishing poorly differentiated stem cell-like colorectal tumours. These data mechanistically and functionally link SRSF3-mediated pri-miRNA processing to hallmark features of cancer.

## Results

### SRSF3 regulates differential processing of polycistronic miRNAs in pluripotent stem cells

We previously generated a reprogrammable tamoxifen inducible *Srsf3* knockout mouse model (*Srsf3*-KO/OKSM) and defined an SRSF3-regulated network of coding RNAs with key roles in pluripotency (17). In addition to protein coding RNAs, abundant SRSF3 binding sites were detected in noncoding RNAs (ncRNAs) both in pluripotent cells and previously in mouse P19 teratocarcinoma cells (Fig. 1*A* and S1*A*) (16, 33) but the functional relevance of SRSF3 binding to ncRNAs in self-renewing cells has not been investigated. The two most enriched ncRNA classes with SRSF3 binding peaks were miRNAs and snoRNAs (Fig. 1*A*). The consensus tetramer binding motif within miRNAs in pluripotent cells was consistent with the previously identified CNNC motif downstream of pri-mRNA stem-loops (Fig. 1*A*) (13). Closer investigation of the SRSF3 miRNA targets demonstrated ample SRSF3 binding within polycistronic miRNA clusters (Table S1), including miR-17-92 that was previously shown to be essential for proliferation and cell-renewal of embryonic stem (ES) cells (30). The overlapping roles of miR-17-92 and SRSF3 in promoting self-renewal (17, 31), as well as previously reported role of SRSF3 in regulating miRNA processing (13, 15) prompted us to further investigate the role of SRSF3 in miR-17-92 processing.

According to previous studies, SRSF3 binding to a CNNC motif within a narrow window of 17-18nt downstream of the Drosha cleavage site is necessary and sufficient to enhance pri-miRNA processing (13, 15). In mouse pluripotent stem cells, SRSF3 binding sites within the miR-17-92 cluster mapped downstream of miR-17 (CNNC at + 18 from Drosha cleavage site), miR-18a (CNNC at + 20), miR-19a (CNNC at + 18) and miR-20a (CNNC at + 18), each overlapping a CNNC motif within this window (Fig. 1*B, C*). Within miR-17a and miR-20a, additional binding peaks with a CNNC motif were detected further downstream (miR-17 CNNC at + 30, miR-20a CNNC at +24 and + 35) (Fig. 1*C*). We also analysed the SRSF3 binding peaks in the two paralog clusters of miR-17-92 (Fig. 1*D*). Two binding peaks were detected within miR-106a-363 (miR-92a-2 CNNC at + 23, miR-363 CNNC at +25) and a single peak within miR-106b-25 (miR-93 CNNC at + 18) (Fig. S1*B*). Of these, miR-93 belongs to the miR-17 family together with miR-17 and miR-20a (Figure 1*D*).

To assess the functional relevance of SRSF3 binding sites within primary miR-17-92 transcript, we measured the levels of pri- and mature miRNAs following SRSF3 depletion by RT-qPCR. The individual mature miRNAs produced from the miR-17-92 pri-miRNA transcript were differently expressed in the control cells (Fig. 1*E* and S1*C*). The loss of SRSF3 did not significantly affect miR-17-92 pri-miRNA abundance (Fig. S1 *D*) but led to a significant decrease in the levels of mature miR-17a and miR-20a with no effect on the other miRNAs produced from the cluster (Fig. 1*E*). We also analysed the expression level of major regulators of the miRNA biogenesis pathway *Drosha, Dicer, Dgcr8, Xpo5* and *Ago2* to confirm that SRSF3 depletion did not affect the expression of the miRNA processing machineries (Fig. S1*E*). These data demonstrate not all SRSF3 binding sites within 17-18 nt distance from the Drosha cleavage sites confer enhanced processing of the upstream miRNAs. This suggests that SRSF3 mediated selective regulation of distinct miRNAs within a single cluster may be determined by factors or sequence features in addition to the previously reported positioning of SRSF3 binding.

### SRSF3 regulated miRNAs control Cdkn1a levels in pluripotent cells

We have previously reported that SRSF3 plays a dual role in mouse pluripotent cells by both enhancing the establishment and maintenance of the pluripotent state and by promoting self-renewal, cell cycle and proliferation (17). The enriched GO terms of differentially expressed genes following SRSF3 depletion in mouse pluripotent cells reflected the functional data, showing enrichment in terms related to cell fate and proliferation (Fig. 2*A*). Our previous work defined mechanisms by which SRSF3 promoted pluripotency (17). Since miR-17-92 derived miRNAs have been reported to promote proliferative and anti-apoptotic effects in various cell types (34–36), we assessed whether SRSF3 controlled cell proliferation during self-renewal through miR-17-92 processing, with the focus on the candidates miR-17 and miR-20a that were affected by SRSF3 depletion (Fig. 1*E*). The cell cycle regulator, cyclin dependant kinase inhibitor *Cdkn1a* (p21) is a *bona fide* target of miR-17a/miR20a (23, 37–39), the 3’UTR of *Cdkn1a* sharing a perfect complementarity with the miR-17/20a seed regions (Fig. 2*B*). In accordance with this, RNA sequencing and RT-qPCR analysis demonstrated increased *Cdkn1a* mRNA expression in SRSF3 depleted mouse pluripotent cells independent of *Trp53* levels (Fig. 2*C, D*). In fact, *Trp53* levels were downregulated, supporting a previous study reporting a p53 independent regulation of p21 in ES cells (40). We also analysed *Trp53* and *Cdkn1a* mRNA expression during the course of reprogramming mouse embryonic fibroblasts (MEFs) into induced pluripotent stem (iPS) cells, which further demonstrated p53 independent regulation of p21 in pluripotent cells. *Cdkn1a* levels followed *Trp53* levels until reprogramming Day 13, the expression diverging as the cells committed to the pluripotent cell fate (Fig. 2*E, S2 A*). *Cdkn1a* expression was affected only in iPS cells where also miRNAs of the miR-17-92 family are abundantly expressed (Fig. 2*E, S2 A-C*) (41). In conclusion, SRSF3 selectively regulates the abundance of miR-17a and miR-20a of the miR-17-92 cluster in pluripotent cells resulting in downregulation of the downstream miRNA target *Cdkn1a*.

**Fig. 2.**
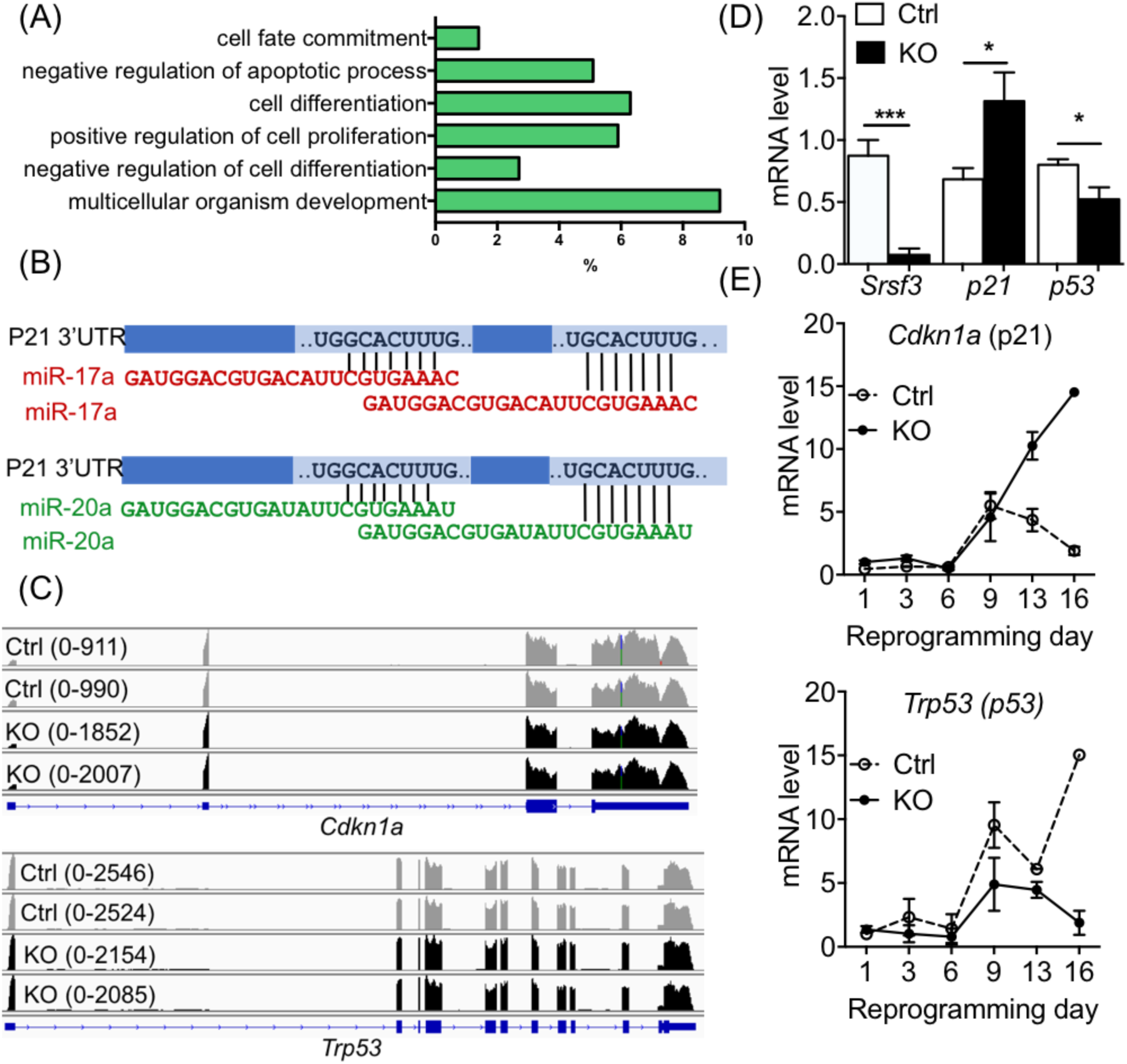
SRSF3 Control *Cdkn1a* levels in pluripotent cells through mIR-17/20a. *(A)* Enriched GO terms of differentially expressed genes following SRSF3 depletion in mouse iPSCs. *(B)* Schematic of miR-17 and miR-20a seed regions that target the *Cdkn1a* 3’UTR at two sites. *(C)* Expression of *Cdkn1a and Trp53* in *Srsf3*-KO and control iPSCs. Window heights are scaled to maximum read counts. *(D)* RT-qPCR quantification of *Srsf3, Cdkn1a* (p21) and *Trp53* (p53) mRNA levels in *Srsf3*-KO and control iPSCs (*p< 0.05, ***<0.0005, Unpaired Student’s t-test, two-tailed, data as mean ± SEM, n=3). *(E)* Quantification of *Cdkn1a* and *Trp53* mRNA expression by RT-qPCR during reprogramming in *Srsf3*-KO and control cells (data as mean ± SEM, n=2). See also Fig. S2.

### Conserved SRSF3-miR-17/20a pathway regulates self-renewal of human colorectal cancer cells

The miR-17-92 cluster including the mature miRNA sequences and their organization is fully conserved in vertebrates (23). The miRNAs produced from the cluster are frequently dysregulated in various cancers, including colorectal cancer, and miR-17-92 was one of the first identified miRNA oncogenes, hence the name OncomiR-1 (23–25, 42). The central role of miR-17-92 in cancer and normal self-renewing cells highlights an intimate link between embryonic development, stem cells and cancer formation. Since SRSF3 is a proto-oncogene able to enhance cell cycle progression and cell proliferation both in normal pluripotent cells and cancer cells, we next investigated whether SRSF3-mediated miR-17-92 played a role in human cancer. We used a Dox-inducible CRISPR/Cas9 system (43)to knockdown SRSF3 and a lentiviral vector encoding SRSF3 and a GFP reporter (SRSF3-T2A-GFP) to overexpress SRSF3 in the human colorectal cancer cell line LIM1215 that endogenously express miR-17-92. Downregulation of SRSF3 in LIM1215 cells led to a selective reduction in miR-17a levels compared to control cells (Fig. 3*A*) while SRSF3 overexpression increased the levels of both miR-17 and its paralog miR-20a (Fig. 3*B*). Similar to mouse pluripotent cells, SRSF3 depletion in LIM1215 cells resulted in increased expression of *CDKN1A*/p21 mRNA and protein without affecting *TP53* levels (Fig. 3*C, D*). The *TP53* mRNA levels were low compared to *CDKN1A* likely explaining very low or non-detectable level of p53 protein in LIM1215 cells (Fig. 3*D*). As predicted based on the increased miR-17/20a levels in SRSF3 overexpressing cells (Fig. 3*B*), CDKN1A/p21 mRNA and protein levels were significantly reduced (Fig 3*E, F*).

**Fig. 3.**
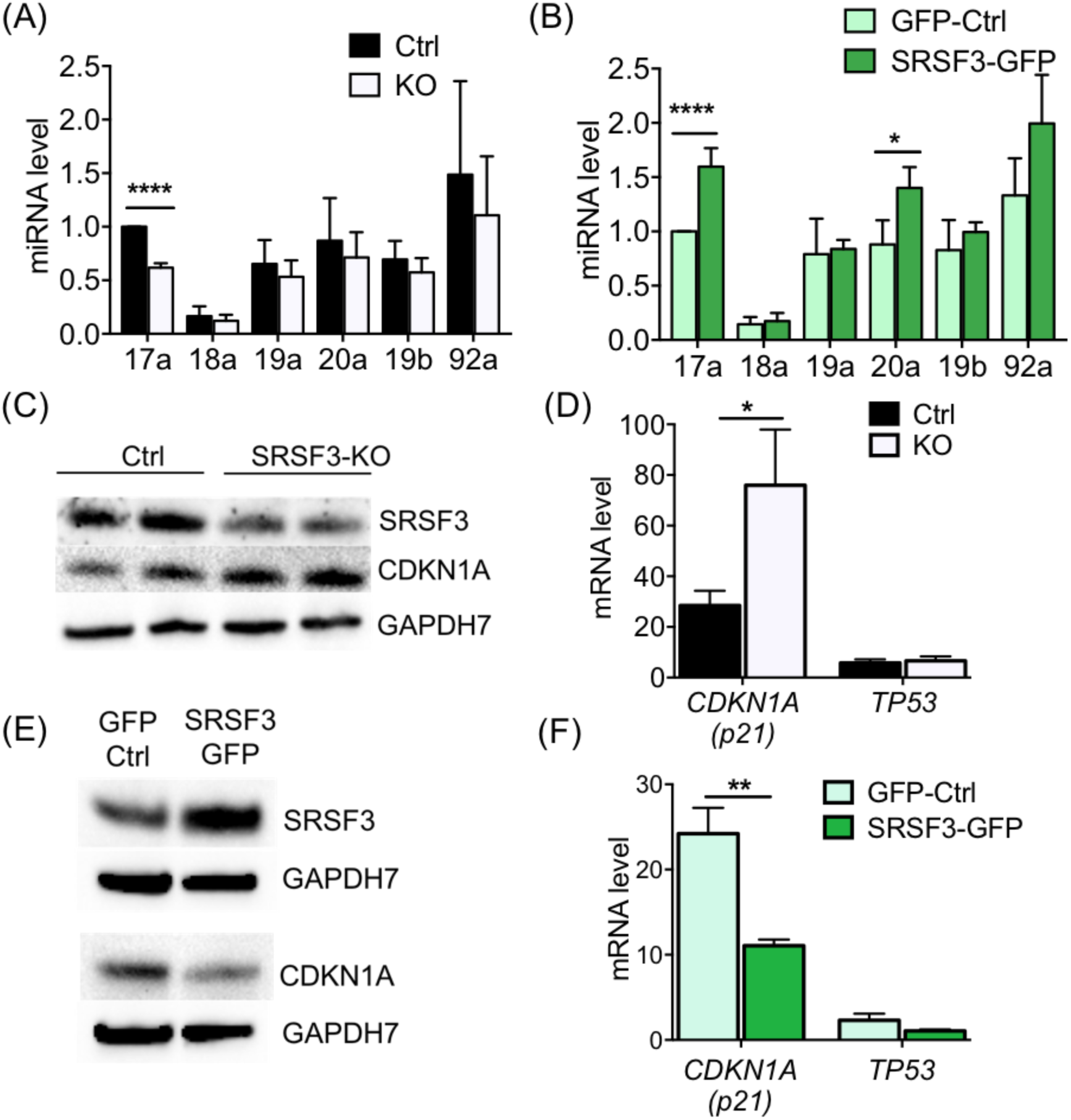
SRSF3-miR-17/20a-Cdkn1a regulation is conserved from mouse pluripotent stem cells to human colon cancer cells. *(A)* TaqMan analysis of the mature miRNAs of miR-17-92 cluster in *Srsf3*-KO and control LIM1215 cells (****p < 0.0001, Unpaired Student’s t-test, two-tailed, data as mean ± SEM, n=3). *(B)* The miRNA TaqMan analysis of the miR-17-92 components in SRSF3-T2A-GFP overexpressing (SRSF3-GFP) and GFP-only control (GFP-Ctrl) LIM1215 cells (* p<0.01, **** p< 0.0001, Unpaired Student’s t-test, two-tailed, data as mean ± SEM, n=4). *(c)* Western blot analysis of SRSF3 and CDKN1A expression in *SRSF3*-KO and control LIM1215 cells. GAPDH7 served as a loading control. *(D)* RT-qPCR quantification of *CDKN1A* (p21) and *TP53* mRNA levels in *SRSF3*-KO and control LIM1215 cells (* p < 0.05, Unpaired Student’s t-test, two-tailed, data as mean ± SEM, n=3). *(E)* Western blot analysis of SRSF3 and CDKN1A expression in SRSF3 overexpressing and control LIM1215 cells. GAPDH7 served as a loading control. *(F)* RT-qPCR quantification of *CDKN1A* (p21) and *TP53* mRNA levels in SRSF3 overexpressing and control LIM1215 cells (* * p < 0.001, Unpaired Student’ s t-test, two-tailed, data as mean ± SEM, n=3).

Next, we examined the cell cycle profile and cell proliferation in LIM1215 cells where SRSF3 levels were modulated. To assess cell proliferation, SRSF3 depleted or overexpressing LIM1215 cells were stained with the cell membrane dye eFluor 670 and analysed by flow cytometry 72 h later. Similar to mouse pluripotent cells and human cell lines (17, 31), SRSF3 overexpression profoundly enhanced the proliferation of LIM1215 cells indicated by the rapid decrease in the intensity of eFluor 670 on the cells while SRSF3 depletion led to a slower proliferation rate as indicated by the slower dilution of the cell surface dye intensity (Fig. 4*A, B* and Fig. S3*A*). The analysis of asynchronous LIM1215 cells demonstrated that high SRSF3 levels resulted in enhanced G1-to-S transition, which was reversed in SRSF3 depleted cells (Fig. 4*C* and Fig. S3*B*). This suggests an early exit from G1 phase following increased SRSF3 expression, which is in line with the enhanced cell proliferation observed above and our data in mouse pluripotent cells. In ES cells the G1 phase is much shorter than somatic cells, lengthening of G1 phase leading to differentiation as seen in SRSF3 depleted pluripotent cells (17, 44–46). These results demonstrate that human colorectal cancer cells fully recapitulate both the molecular and cellular phenotype of mouse pluripotent stem cells and highlight the functional relevance of the conserved SRSF3-mediated regulation of self-renewal through miR-17a/miR20a and p21.

**Figure 4.**
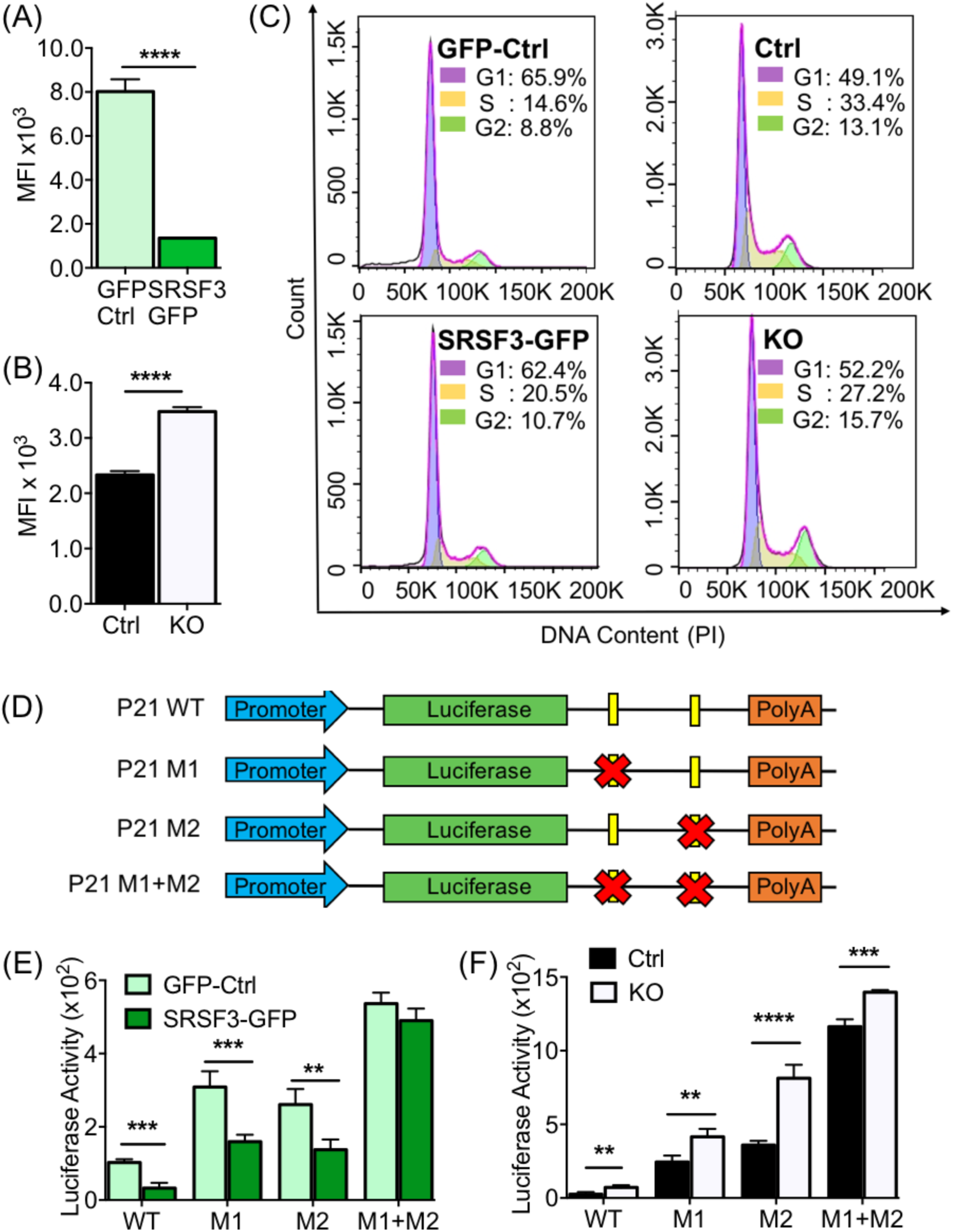
SRSF3-miR17/20a-CDKN1A pathway regulates self-renewal of human colorectal cancer cells. *(A)* Quantification of Mean Fluorescent Intensity (MFI) of eFluor 670 labelled SRSF3 overexpressing and control LIM1215 cells (****p < 0.0001, Unpaired Student’s t-test, two-tailed, data as mean ± SEM, n=3). *(B)* Quantification of Mean Fluorescent Intensity (MFI) of eFluor 670 labelled *SRSF3*-KO and control LIM1215 cells (***p<0.0005, Unpaired Student’s t-test, two-tailed, data as mean ± SEM, n=3). *(C)* DNA histograms of PI stained LIM1215 cells as analysed by flow cytometry. Histograms were generated using cell cycle analysis tool in FlowJo software. Percent cells in each phase is indicate in the panels. *(D)* Schematic of the Firefly luciferase reporter constructs used (Cite Wang et al Paper) *(E)* Luciferase activity in SRSF3 overexpressing and control LIM1215 cells transfected with each reporter construct (***p<0.0005, **P < 0.005, Unpaired Student’s t-test, two-tailed, data as mean ± SEM, n=4). *(F)* Quantification of luciferase activity in *SRSF3*-KO and control LIM1215 cells transfected with each reporter construct (**p< 0.005, ***p<0.0001****P<0.0001, Unpaired Student’s t-test, two-tailed, data as mean ± SEM, n=4). See also Fig. S3.

### *CDKN1A is the main downstream target of* miR-17/20a *in self-renewing cells*

Individual miRNAs often target multiple components of a pathway. CDKN1A belongs to the CyclinE/CDK2 pathways consisting of multiple cell cycle inhibitors. TargetScan (39) predicted miR-17/20a binding to the 3’UTR of *Cdkn1a, Rbl1, Rbl2* and *Rb1*. Surprisingly, analysis of RNA sequencing data of SRSF3 knockout and control mouse iPS cells (17) revealed that only *Cdkn1a* was significantly affected by the SRSF3 depletion (Fig. S3*C*). We also analysed predicted miR-17/20a target genes associated with cell cycle regulation and supported by miRPathDB (47). Among the 16 genes analysed, only *Cdkn1a* expression was increased more than 1.5-fold in SRSF3 depleted cells (Table S2). These data suggest that the cell proliferation and cell cycle phenotypes observed in SRSF3 modulated mouse pluripotent and potentially human cancer cells are largely mediated through miR-17a/20a targeting *Cdkn1a*. To address the direct link between SRSF3, miR-17/20a and *Cdkn1a*, we used luciferase reporters containing either the wildtype *Cdkn1a* 3’UTR downstream of a constitutively expressed firefly luciferase or the 3’UTR where one or both of the miR-17/20a target sites are mutated (Fig. 4*D*; M1, M2 or M1+M2)(20). Luciferase activity was measured in control, SRSF3 depleted or overexpressing LIM1215 cells. Following GFP expression (control), the luminescence signal was increased when either one of the two binding sites were mutated and mutation of both sites led to further signal increase indicative of an independent and additive effect of the two miRNA target sites on Cdkn1a expression (Fig. 4*E*, light green bars). SRSF3 overexpression resulted in a significantly reduced luciferase activity when the cells expressed luciferase with WT, M1 or M2 3’UTRs which was abolished in cells expressing the M1+M2 construct (Fig. 4*E*, dark green bars and Fig. S3*D*). Similar to molecular and cellular analysis, SRSF3 depletion reversed the effect, verifying the direct role of SRSF3 on CDKN1A/p21 expression via miR-17/20a (Fig. 4*F*, Fig. S3*E*).

### SRSF3-miR-17/20a-CDKN1A pathway as a predictive marker for advanced colorectal cancer

Dysregulation of SRSF3 and miR-17-92 cluster miRNAs has been reported in colorectal cancer (48-50). Analysis of RNA isolated from the tumour and adjacent normal tissue of 25 colorectal cancer patients not subjected to chemo-or radiotherapy demonstrated significantly increased SRSF3 levels in the tumour compared to their paired normal tissue (Fig. 5*A* and Fig. S4*A*). As predicted by the LIM1215 data (Fig. 3), *CDKN1A* levels were significantly reduced in tumour samples with no difference in *TP53* levels (Fig. 5*B*). Importantly, miRNAs of the miR-17-92 cluster were significantly upregulated in tumour samples with particularly high levels of miR-17 and miR-20a, suggesting that the *SRSF3-*miR-17/20a*-CDKN1A* pathway operates in colorectal cancer (Fig. 5*C*). Looking at individual patients, 60% of patients (15 / 25) displayed the SRSF3 signature, suggesting that this pathway may be active in a large fraction of CRC patients (Fig. 5*D* and Table S3).

**Figure 5.**
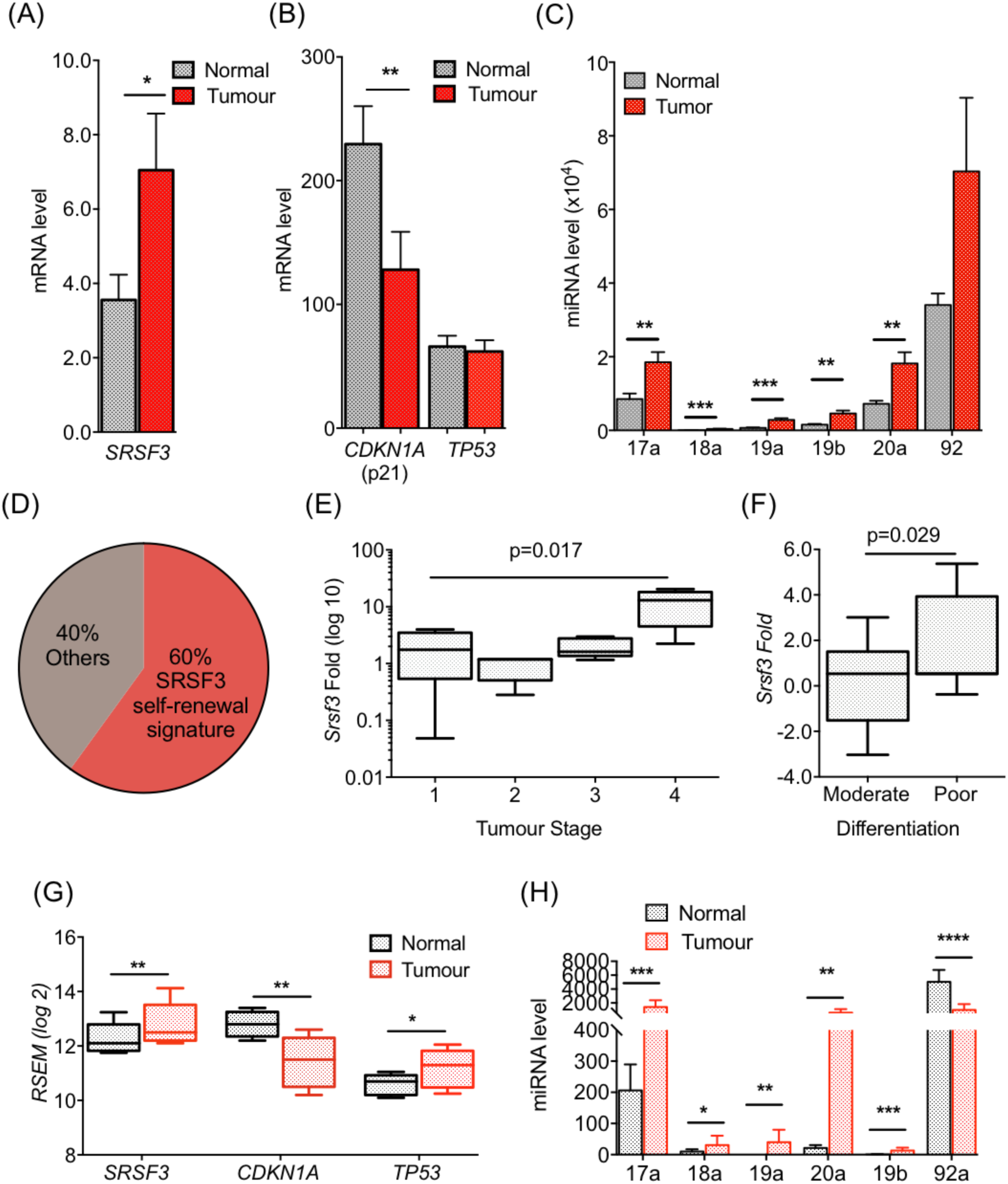
SRSF3 self-renewal signature is a predictive marker for advanced colorectal cancer. *(A)* RT-qPCR quantification of *SRSF3* mRNA levels in colorectal tumours and paired normal samples (*p<0.05, Unpaired Student’s t-test, two-tailed, data as mean ± SEM, n=25). *(B)* RT-qPCR quantification of *CDKN1A* and *TP53* mRNA levels in colorectal tumours and paired normal samples (*p<0.05, Unpaired Student’s t-test, two-tailed, data as mean ± SEM, n=25). *(C)*TaqMan analysis of the mature miRNAs of miR-17-92 cluster in colorectal tumours and paired normal samples (**p<0.005, ***p<0.0005, Unpaired Student’s t-test, two-tailed, data as mean ± SEM, n=25). *(D)* Quantification of the fraction of patients with SRSF3 self-renewal signature. *(E)* First, 2^nd^ (median), and 3^rd^ quartiles (box) and ±95% whiskers of *SRSF3* expression in T1 (n=3), T2 (n=5), T3 (n=11) and T4 (n=3) tumours. *(F)* First, 2^nd^ (median), and 3^rd^ quartiles (box) and ±95% whiskers of SRSF3 expression in patients with moderately differentiated tumours (n=7) and patients with poorly differentiated tumours (n=7). *(G)* First, 2^nd^ (median), and 3^rd^ quartiles (box) and ±95% whiskers of *SRSF3, CDKN1A* and *TP53* expression in TCGA COAD data (version 2016_01_28, tumour n=459 and normal n=41) *p<0.05, **p<0.005 Unpaired Student’s t-test, two-tailed, data as mean ± SEM). *(H)* Relative mRNA expression of mature miRNAs of miR-17-92 cluster in TCGA COAD data (version 2016_01_28, tumour n=272 and normal n=8) (*p< 0.05, **p<0.005, ***p<0.0005 ****p< 0.0001, Unpaired Student’s t-test, two-tailed, data as mean ± SEM).

We then analysed the relevance of the *SRSF3-*miR-17/20a*-CDKN1A* pathway for clinical characteristics including Overall stage, Grade, T-stage, N-stage, M-stage, Gender and Age. SRSF3 expression was significantly associated with T4 stage (advanced cancer with tumour penetrating through all layers of colon and possibly into adjacent organs) compared to T1 stage tumours (p=0.002) (Fig. 5*E*). Strikingly, SRFS3 expression was significantly associated with poorly differentiated stem cell like tumours (high grade/anaplastic) compared to moderately differentiated (intermediate grade) tumours (p=0.029) (Fig. 5*F*), in agreement with our data on the role of SRSF3 in self-renewal (17). Taken together these results suggest that SRSF3-regulated OncomiR-1 processing may be a marker for advanced/higher grade colorectal cancer with cancer stem cell characteristics.

Due to the relatively small size of our initial patient cohort, we also analysed the TCGA COAD data (version 2016_01_28) for the relevance of the *SRSF3-*miR-17/20a*-CDKN1A* pathway. The TCGA data set included mRNA expression data for 459 tumour and 41 normal samples as well as miRNA expression data for 272 tumour samples and 8 normal samples. The TCGA data supported the analysis of our initial patient cohort with an increase in SRSF3 expression (fold-change 1.3), downregulation of *CDKN1A* levels (fold change 0.4) and upregulation of miRNAs of the miR-17-92 cluster (miR-17a, 18a, 19a, 20a and 19b) in tumour samples compared to normal tissue (Fig. 5*G, H*). Among the OncomiR-1 miRNAs, the most dramatic increase was detected in miR-17a and miR-20a levels (Fig. S4*B*). While no difference in *TP53* levels was observed in our patient cohort (Fig. 5*B*), in the larger TCGA cohort *TP53* levels were increased (Fig. 5*G*). This aligns with our data in pluripotent cells and confirms the p53-independent regulation of p21 also in colorectal cancer patients. In conclusion, SRSF3 acts as an upstream regulator of p21 expression through miR-17a and miR20a both in normal self-renewing cells and in the context of colorectal cancer.

## Discussion

MicroRNA biogenesis needs to be tightly regulated to maintain tissue specific miRNA expression patterns. In addition to transcriptional control, RBPs can recognize sequences within pri-miRNAs and modulate their processing post-transcriptionally. In particular, strikingly different levels of individual mature miRNAs derived from polycistronic pri-miRNA transcripts cannot be explained by transcriptional regulation. The differential regulation of individual miRNAs derived from a polycistron does not only affect normal development and homeostasis but dysregulation of individual miRNAs can play a role in tumourigenesis (38). Here we demonstrate that SRSF3 specifically regulates the processing of two paralog miRNAs from the miR17-92/OncomiR-1 polycistron, which leads to enhanced cell cycle progression and cell proliferation, and is a hallmark of poorly differentiated high-grade tumours. This conserved mechanism of selective miRNA processing demonstrates how post-transcriptional regulation of miR17-92/OncomiR-1 processing plays a central role in tumour promoting properties and how RBPs such as SRSF3 can define the functional output of polycistronic pri-miRNAs.

The SRSF3 consensus binding site CNNC and/or experimentally defined SRSF3 binding sites are found downstream of Drosha cleavage site in the majority of mono- and polycistronic pri-miRNAs (13). The CNNC motif and SRSF3 binding has been shown to enhance the processing of pri-miRNAs but the presence of the motifs appear to affect individual miRNAs differentially (13), suggesting that not all CNNC sites are functionally identical or that some sites are only used in distinct cellular conditions. Our data shows that although the CNNC motif and SRSF3 binding sites are found in four miRNAs within the polycistronic miR17-92 at the ‘ideal’ 17-18nt distance from the Drosha cleavage site, only the processing of miR-17 and miR-20a is modulated by SRSF3 in mouse pluripotent cells and human colorectal cancer cells. The distinguishing feature downstream of miR-17 and miR-20a was the presence of an additional CNNC site further downstream of the CNNC within the 17-18nt window from Drosha cleavage site. This suggests that although a single SRSF3 binding site may be sufficient *in vitro* or within monocistronic miRNAs, in polycistronic miRNAs the synergistic interaction between neighbouring SRSF3 binding sites may be required for enhanced processing by Drosha at the apical junction.

Functional dissection of miR-17-92 indicates that individual miRNAs of the cluster regulate distinct pathways during development and disease and the biological effects of miR-17-92 heavily depend on cellular context (38). For example, miR-17-92 can promote angiogenesis in tumour cells through the activity of miR-18 and miR-19 (51) but repress it in endothelial cells through miR-17 (52). This complexity conferred by multiple components of the polycistronic clusters highlights the contribution of individual components of miR-17-92 cluster to the overall oncogenic activity under different biological contexts. Similarly, miR-17-92/OncomiR-1 has been implicated as both an oncogene and tumour suppressor depending on the relative abundance of individual miR-17-92 miRNAs and biological context (42, 53). Strikingly, the two miRNAs regulated by SRSF3 within the miR-17-92 cluster are paralogs targeting largely overlapping mRNAs. Among the targets of miR-17 and miR-20a is the cell cycle inhibitor Cdkn1a/p21 that enhances G1-to-S transition in the self-renewal of both normal and malignant cells (54). Our data shows how SRSF3 specifically enhances the processing of miR-17 and miR-20a without affecting other miRNAs within the same cluster in both mouse pluripotent cells and human LIM1215 CRC cells. In both species, modulation of SRSF3 leads to altered Cdkn1a/p21 expression independent of its canonical upstream regulator p53. Accordingly, cell cycle progression and cell proliferation were promoted by SRSF3 overexpression and inhibited by its downregulation, linking SRSF3 to Cdkn1a/p21 and cell cycle. Previous work has implicated SRSF3 in cell proliferation and cell cycle (17, 31, 50) but these reports have not provided conclusive evidence for the role of SRSF3 in cell cycle progression. Intriguingly, the expression of SRSF3 is regulated during cell cycle, its expression induced in late G1 or early S (55). The promoter of SRSF3 contains two consensus binding sites for E2F, a transcription factor involved in regulating miR17-92 and the cell cycle (55, 56). This suggests that the cell cycle phenotype regulated by SRSF3 and miR17/20a is linked in self-renewing and cancer cells. While a shorter G1 phase is critical during early embryogenesis, an inappropriately shorter G1 phase in more differentiated cells can be associated with uncontrolled cell growth leading to tumour formation (20, 57).

The analysis of tumour normal pairs from a CRC patient cohort revealed increased expression of *SRSF3*, miR-17 and miR-20a with a corresponding decrease in *CDKN1A* expression without a change in *TP53* levels which was further supported by TCGA data set (~400 tumours). Strikingly, this ‘signature’ was sufficient to classify patients with poorly differentiated tumours. While we do not have the survival information of the patients used for this study, the high grade or poorly differentiated histology is known to be associated with poor patient survival (58–60). SRSF3 protein expression is upregulated in putative CD133+ colon cancer stem cells in comparison to CD133-cells and its depletion slowed down cell proliferation of Caco-2 colorectal cancer cells, supporting the role of SRSF3 in tumorigenicity of colon cancer (61). A single G-to-A mutation in the terminal loop of pri-mir-30c-1 in breast and gastric cancer patients has been shown to alter the secondary pri-miRNA structure facilitating SRSF3 binding and processing by Drosha (62). Similarly, a C-to-T mutation within the CNNC motif downstream of pri-miR-16 has been associated with chronic lymphocytic leukemic (63). Here we demonstrate that SRSF3 overexpression alone is sufficient to favour the production of oncogenic miRNAs from the miR-17-92/OncomiR-1 cluster, leading to rapid cell cycle progression and proliferation in colorectal cancer. These data underline the central role of SRSF3-mediated pri-miRNA in tumorigenesis, suggesting that concurrent dysregulation of SRSF3 and oncogenic miRNAs may be one of the hallmark features of cancer cells.

## Methods

### Generation of iPS/ES cell lines and reprogramming

iPS cell lines with induced SRSF3 knock-down was generated from *Srsf3*^tm1a(KOMP)Mbp^ knockout-first mice (Knock Out Mouse Project, KOMP, UC Davis Mouse Biology Program) via Flpe recombination as described before (17). SRSF3-BAC ES cells and EGFP-NLS control cells were generated as previously by transfecting BAC-transgene encoding mouse *Srsf3* C-terminally tagged with EGFP or EGFP-NLS vector into mouse JM8A3.N1 ES cells (64). Transfected cells were selected using geneticin, followed by enrichment for EGFP-positive cells by FACS. Reprogramming was performed using conditional SRSF3 knock-out and control MEFs as described before (17). All animal works were performed according to the Australian code for the care and use of animals for scientific purposes (NHMRC) and approved by the Monash University Animal Ethics Committee.

### Generation of LIM1215 cell lines and general cell culture

The LIM1215 cells were cultured at 37°C under 5% CO_2_. The cells were maintained in RPMI supplemented with 10% FBS, penicillin/streptomycin and Glutamax. The SRSF3 overexpressing cells were generated using a lentiviral overexpression construct carrying a cDNA encoding human *SRSF3* followed by *T2A* self-cleaving peptide and *EGFP* (*SRSF3-T2A-EGFP*). A vector carrying only *EGFP* was used as a control. The SRSF3 knock-out LIM1215 cells were generated using an inducible sgRNA construct targeting exon 2 of human *SRSF3* gene (43). The forward and reverse strand oligonucleotides were annealed and ligated into pFH1tUTG-T2A-GFP plasmid. The pFH1tUTG-Cas9-T2A-mCherry plasmid was received from Marco J Herold at the Walter and Eliza Hall Institute of Medical Research, Australia. Both constructs were packaged in to lentiviral vectors as described before (43). The LIM1215 cells were co-infected with both constructs and the cells that were double positive for both GFP and mCherry expression were sorted using flow-cytometry. The SRSF3 knock-down was induced with Doxycycline at a final concentration of 2μg/ml.

### RNA extraction and quantitative real-time PCR

Total RNA was isolated using the TRI Reagent (Sigma-Aldrich) and subjected to DNaseI treatment. RNA was used for cDNA synthesis with SuperScript III Reverse Transcriptase (Thermo Scientific). Quantitative real-time PCR was performed in an AB 7500 real-time PCR system (Applied Biosystems) with Luminaris HiGreen qPCR Master Mix-low ROX (Thermo Scientific) and 0.3uM primer mix (Table S4). The cycle threshold value (CT) was calculated for each sample and normalised to *Hprt*. The relative mRNA levels were calculated using ΔΔCT method.

For mature miRNA analysis, Applied Biosystems TaqMan MicroRNA Assays were used. cDNA was generated using the Applied Biosystems® TaqMan® MicroRNA Reverse Transcription Kit according to the manufacturer’s instructions. Resulting cDNA was amplified with the SensiFAST Probe Hi-ROX Kit (Bioline). The fold change in expression were calculated by the ΔΔCT method using snoRNA202 and U6 RNA as reference small RNAs in mouse and human cells, respectively.

### Western Blotting

For all Western blots, cells were lysed in RIPA buffer (50mM Tris-HCl pH 8, 150mM NaCl, 1% Nonidet P40, 0.5% Sodium deoxycholate, 0.1% SDS). NuPAGE 4–12% gradient Bis-Tris gel system (Thermo Scientific) was used in reducing conditions. The nitrocellulose membranes were probed with antibodies against SRSF3 (Sigma-Aldrich), CDKN1A (Cell Signalling Technologies), and GAPDH7 (Cell Signalling Technologies), followed by HRP-conjugated secondary antibodies (Biorad). The blots were developed using Amersham ECL Western Blotting Detection Reagents (GE Healthcare) and visualized with the Biorad ChemiDoc MP Imaging System. The band intensities were quantified with ImageJ.

### Cell Proliferation Assay

SRSF3 over expressing, SRSF3 depleted and control LIM1215 cells were stained with the cell proliferation dye eFluor670 (10μM). 1.5×10^5^ cells each were plated on a 12-well plate with 3 replicates per cell line. Cells were analysed using the flow-cytometry after 72hrs. DAPI was used as a viability marker.

### Cell Cycle Analysis

Cell cycle distribution was determined by flow cytometry. SRSF3 over expressing, SRSF3 depleted and control LIM1215 cells were collected and fixed in 70% ethanol. The fixed cells were washed with PBS, treated with RNase (100μg/ml) and propidium iodide (50μg/ml) and analysed by flow cytometry.

### Luciferase Assay

The luciferase constructs pGL-p21UTR WT, M1, M1, M1 + M2 was a gift from Robert Blelloch. LIM1215 cells overexpressing SRSF3, depleted of SRSF3 as well as their respective controls were transfected with 4μg of each construct using Lipofectamin 2000 reagent. After 48hrs the cells were collected with trypsin and 1x 10^5^ cells each was used in triplicates. The assay was conducted using the Pierce Firefly Luciferase Glow Assay Kit (Thermo Scientific) according to the manufacturer’s instructions.

### Patient Samples

RNA samples from tumour and their paired normal tissues of 25 CRC patients were received from the Cabrini Hospital, Malvern, Australia. Twenty five patients with primary adenocarcinomas (n = 21), adenocarcinoma mucinous (n=2), adenocarcinoma signet (n=1) or adenomas (n = 1) underwent surgery in 2012/2013 at Cabrini Hospital (Malvern, Victoria). The study was approved by the Cabrini Human Research Ethics Committee (ethics no, 05-11-04-11) and the Monash Human Research Ethics committee (ethics no. 9731). All patients provided written informed consent. Tumour specimens and matched normal colon tissues were freshly stored in RNAlater (Qiagen) for RNA extraction.

## Acknowledgements

This work was supported by the Victorian Government Operational Infrastructure Support Scheme and National Health and Medical Research Council (NHMRC) grants to MLA (GNT1043092 and GNT1138870) as well as by NHMRC grants to HEA and PJM (GNT1010429 and GNT1100531) and in part by “Let’s Beat Bowel Cancer” (www.letsbeatbowelcancer.com), a benevolent fund raising and public awareness foundation. The funders had no part in the design, conduct, outcomes, decision to publish or the drafting of this manuscript. The work was supported by Monash University and Hudson Institute of Medical Research technology platforms FlowCore, Monash Animal Research Platform and MHTP Genomics Facility.

