## Supplemental Information for "SRSF3 confers selective processing of miR-17-92 cluster to promote tumorigenic properties in colorectal cancer"

### Supplementary Information

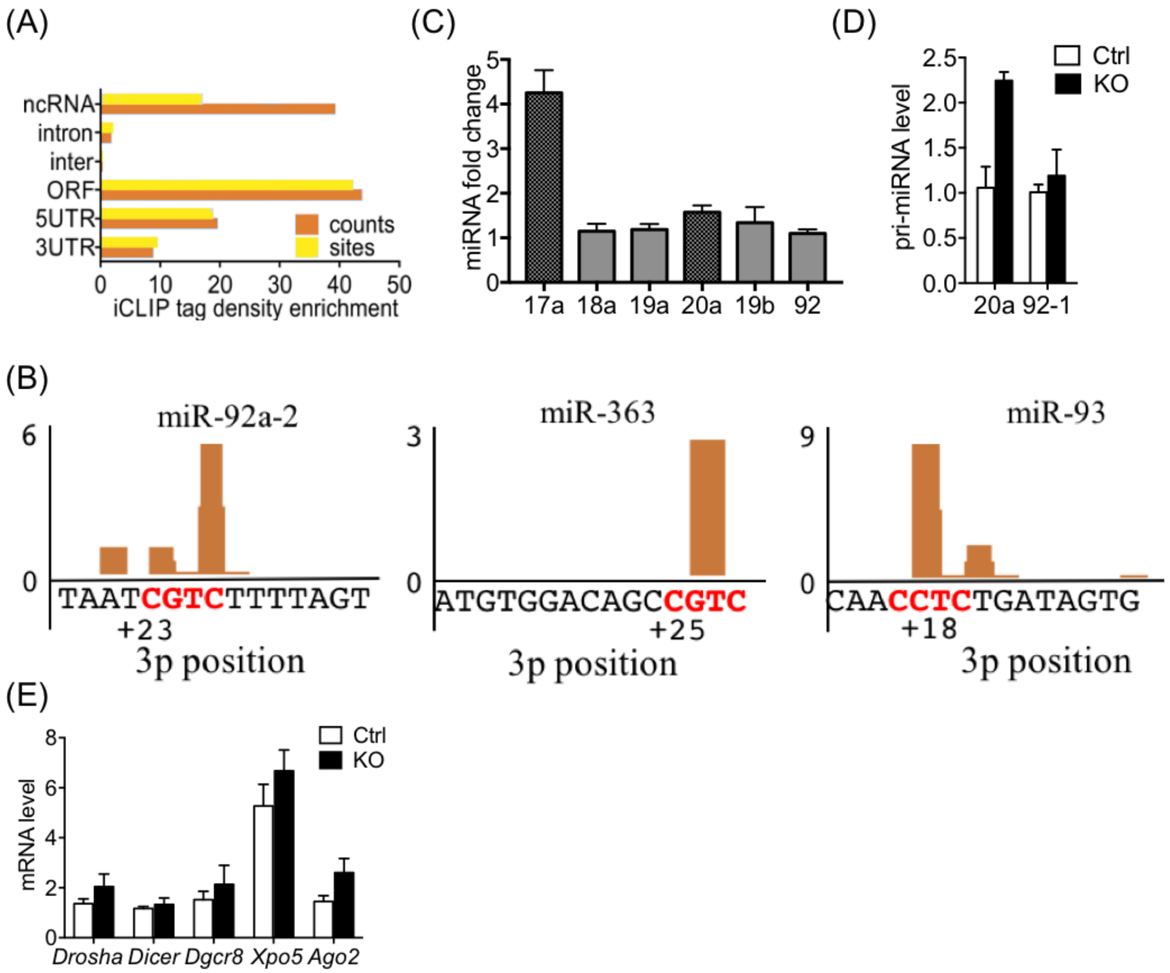


**Fig. S1**. (*A*) Distribution of significant SRSF3 crosslink sites (FDR<0.05) over different transcript regions normalised to feature length. (*B*) SRSF3 iCLIP binding peaks within the components of paralogue clusters miR-106a-363 and miR-106b-25 in mouse ES cells. The 3P position is counted from the 3' Drosha cleavage site as described in [1]. The CNNC motifs are marked in red colour. (*C*) Fold changes of the miR-17-92 cluster miRNAs measured in Figure 1*E*. (*D*) RT-qPCR quantification of pri-miRNA expression in *Srsf3*-KO and control iPSCs. Two different primer pairs around miR-20a and miR-92-1 stem loop regions were used. ($p>0.05$, Unpaired Student's t-test, two-tailed, data as mean ± SEM, n=3). (*E*) RT-qPCR quantification of *Drosha, Dicer,*

*Dgcr8, Xpo5 and Ago2* mRNA levels in *Srsf3*-KO and control iPSCs ($p>0.05$ for all genes analysed, Unpaired Student's t-test, two-tailed, data as mean ± SEM, n=3).

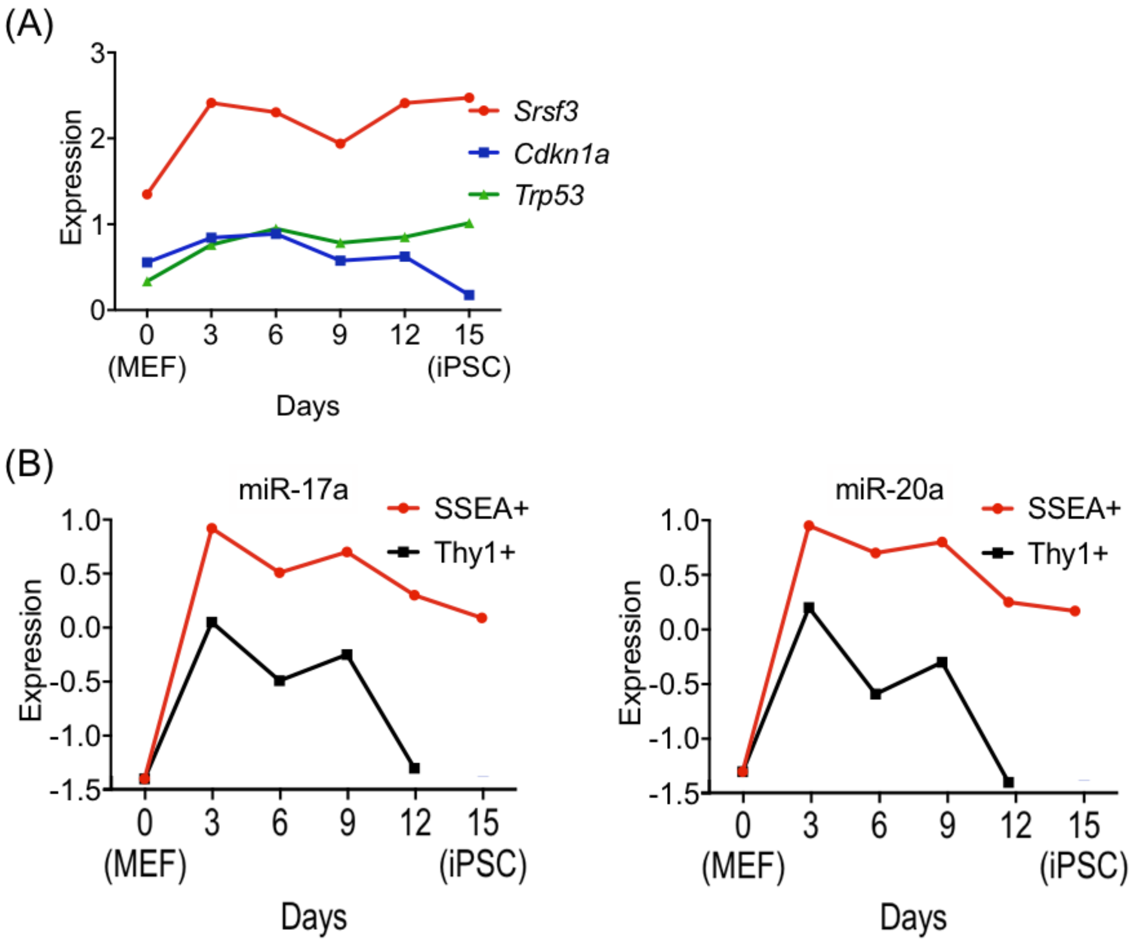


**Fig. S2.** (*A*) *Srsf3, Cdkn1a and Trp53* mRNA expression during reprogramming in the SSEA1+ population. The graph is based on data from (Polo 2012 paper reference). (*B*) Comparison of miR-17 and miR-20a expression in SSEA1+ and Thy1+ populations during reprogramming. The graphs are based on data from [2].

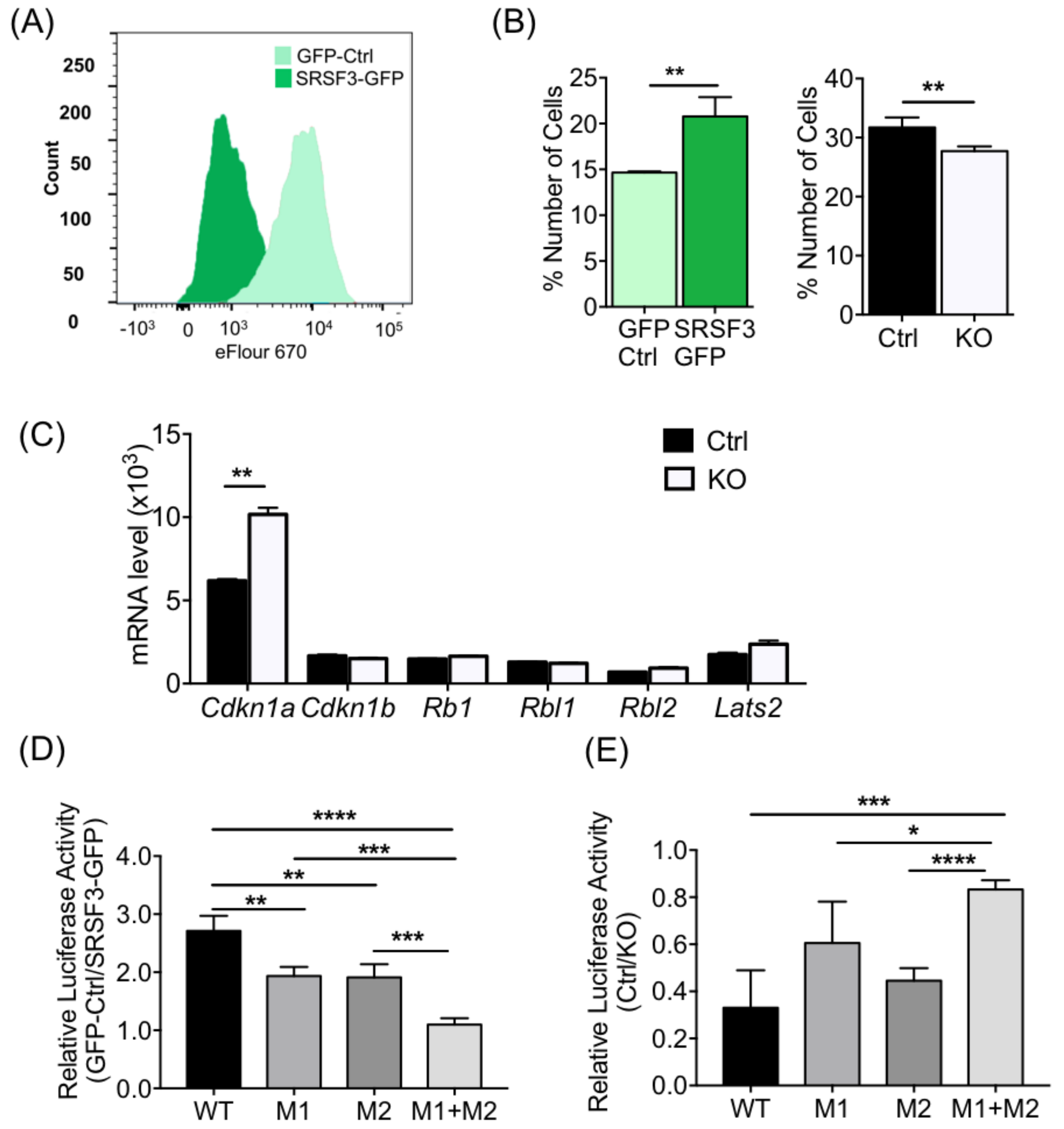


**Fig. S3.** (*A*) Representative histogram of eFluor 670 intensity in SRSF3 overexpressing and control LIM1215 cells. (*B*) Quantification of percent number of cells in S phase in SRSF3 overexpressing and SRSF3-KO cells relative to their respective controls (**$p<0.005$, Unpaired Student's t-test, two-tailed, data as mean ± SEM, n=4). (*C*) Relative mRNA expression of CyclinE/CDK2 inhibitors, *Cdkn1a, Cdkn1b, Rb1, Rbl1, Rbl2* and *Lats2* in SRSF3-KO and control iPSCs (**$p<0.005$, Unpaired Student's t-test, two-tailed, data as mean ± SEM, n=2). (*D*) Ratio of relative luciferase activity between control and SRSF3 overexpressing LIM1215 cells following transfection with luciferase reporters (**p<0.005, ***p< 0.0005, ****p<0.0001, Unpaired Student's t-test, two-tailed, data as mean ± SEM, n=4). (*E*) Ratio of relative luciferase activity between control and SRSF3-KO LIM1215 cells following transfection with luciferase reporters (**p<0.005, ***p< 0.0005, ****p<0.0001, Unpaired Student's t-test, two-tailed, data as mean ± SEM, n=4).

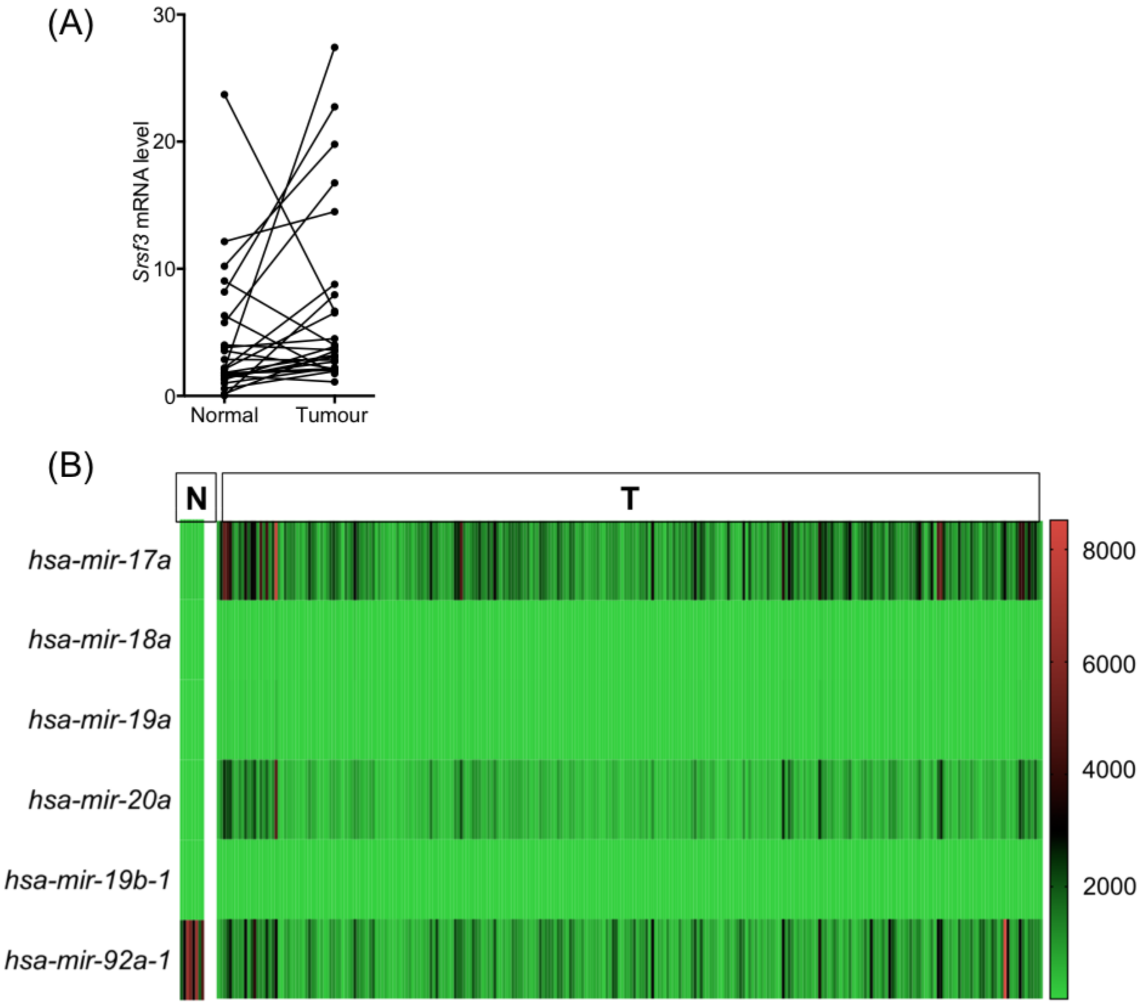


**Fig. S4.** (A) RT-qPCR quantification of SRSF3 mRNA levels in colorectal tumours and paired normal samples presented as individual pairs (*p<0.05, Unpaired Student's t-test, two-tailed, data as mean ± SEM, n=25). (B) Heat map for mRNA expression of miR-17-92 components in TCGA COAD data (version 2016_01_28) of tumour (n=272) and normal (n=8) samples.

**Table S1. SRSF3 binding peaks within miRNA.**

| Chromosome | miRNA | Category | Total sites | Site density |
|---|---|---|---|---|
| chr12 | Mir5099 | monicistronic miRNA | 50 | 0.675676 |
| chr17 | Mir5125 | monicistronic miRNA | 46 | 0.582278 |
| chr9 | mmu-mir-6236 | monicistronic miRNA | 44 | 0.357724 |
| chr14 | Mir20a | polycistronic miRNA | 43 | 0.401869 |
| chr7 | Mir293 | polycistronic miRNA | 35 | 0.4375 |
| chr7 | Gm22133 | unannotated miRNA | 32 | 0.293578 |
| chr2 | Gm24049 | unannotated miRNA | 28 | 0.28 |
| chr7 | Mir291a | polycistronic miRNA | 20 | 0.243902 |
| chr9 | Gm26377 | unannotated miRNA | 15 | 0.138889 |
| chr12 | Mir1193 | polycistronic miRNA | 11 | 0.0909091 |
| chr14 | Mir92-1 | polycistronic miRNA | 9 | 0.1125 |
| chr12 | Mir1197 | polycistronic miRNA | 7 | 0.0583333 |
| chr17 | mmu-mir-8094 | polycistronic miRNA | 7 | 0.0598291 |
| chr12 | Mir654 | polycistronic miRNA | 6 | 0.0714286 |
| chr13 | Mirlet7a-1 | polycistronic miRNA | 6 | 0.06 |
| chr19 | Mir5114 | monicistronic miRNA | 2 | 0.0327869 |
| chr2 | mmu-mir-7005 | monicistronic miRNA | 2 | 0.0289855 |
| chr11 | Gm25994 | unannotated miRNA | 1 | 0.00990099 |

**Table S2. Gene expression of miR-17/20a targets associated with cell cycle regulation.**

| Gene ID | Gene Name | Fold-change (Ctrl vs KO) |
|---|---|---|
| ENSMUSG00000070348 | *Ccnd1* | 0.85 |
| ENSMUSG00000000184 | *Ccnd2* | 1.42 |
| **ENSMUSG00000023067** | ***Cdkn1a*** | **1.65** |
| ENSMUSG00000027490 | *E2f1* | 0.89 |
| ENSMUSG00000022105 | *Rb1* | 1.11 |
| ENSMUSG00000027641 | *Rbl1* | 0.94 |
| ENSMUSG00000057329 | *Bcl2* | 1.49 |
| ENSMUSG00000052934 | *Fbxo31* | 0.95 |
| ENSMUSG00000020184 | *Mdm2* | 1.34 |
| ENSMUSG00000034462 | *Pkd2* | 1.19 |
| ENSMUSG00000013663 | *Pten* | 1.01 |
| ENSMUSG00000020167 | *Tcf3* | 0.84 |
| ENSMUSG00000024515 | *Smad4* | 1.10 |
| ENSMUSG00000031016 | *Wee1* | 0.79 |
| ENSMUSG00000022346 | *Myc* | 0.93 |
| ENSMUSG00000031666 | *Rbl2* | 1.33 |

**Table S3. Expression of *SRSF3*, miR-17/20a and *CDKN1A* in CRC tumours compared to their paired normal samples.**

| Patient # | *SRSF3* | *CDKN1A* | miR-17 | miR-20a |
|---|---|---|---|---|
| 1 | ↑ | ↓ | ↑ | ↑ |
| 2 | ↑ | ↓ | ↑ | ↑ |
| 3 | ↓ | ↓ | ↓ | ↑ |
| 4 | ↑ | ↓ | ↑ | ↑ |
| 5 | ↑ | ↓ | ↑ | ↑ |
| 6 | ↓ | ↑ | ↑ | ↑ |
| 7 | ↑ | ↓ | ↑ | ↑ |
| 8 | ↑ | ↓ | ↑ | ↑ |
| 9 | ↑ | ↓ | ↑ | ↑ |
| 10 | ↑ | ↓ | ↑ | ↑ |
| 11 | ↑ | ↓ | ↑ | ↑ |
| 12 | ↓ | ↓ | ↑ | ↑ |
| 13 | ↓ | ↓ | ↓ | ↑ |
| 14 | ↑ | ↑ | ↑ | ↑ |
| 15 | ↑ | ↓ | ↑ | ↑ |
| 16 | ↑ | ↓ | ↑ | ↑ |
| 17 | ↑ | ↓ | ↑ | ↑ |
| 18 | ↓ | ↓ | ↑ | ↑ |
| 19 | ↑ | ↓ | ↑ | ↑ |
| 20 | ↓ | ↑ | ↑ | ↓ |
| 21 | ↓ | ↓ | ↓ | ↓ |
| 22 | ↑ | ↑ | ↓ | ↓ |
| 23 | ↑ | ↑ | ↑ | ↑ |
| 24 | ↑ | ↓ | ↓ | ↑ |
| 25 | ↑ | ↓ | ↑ | ↓ |

**Table S4. Primers used in this study (F= forward primer, R= reverse primer).**

| Gene | | Sequence |
|---|---|---|
| *Srsf3* (mouse) | F | TGAATTAGAACGGGCTTTTGG |
| | R | TTCACCATTCGACAGTTCCAC |
| *Hprt* (mouse) | F | TGTTGTTGGATATGC |
| | R | TGCGCTCATCTTAGG |
| Pri-miRNA 92-1 (mouse) | F | TCTGCTGTGCAAATCCATGC |
| | R | TCTTCTGGTCACAATCCCCAC |
| Pri-miRNA 20a (mouse) | F | CAAAACTGATGGTGGCCTGC |
| | R | GCTCAATAACAGGACAGTTGGC |
| *Cdkn1a*/p21 (mouse) | F | TTGCACTCTGGTGTCTGAGC |
| | R | TGCGCTTGGAGTGATAGAAA |
| *Trp53*/p53 (mouse) | F | CCCGAGTATCTGGAAGACAGG |
| | R | GTAAGGATAGGTCGGCGGTTC |
| *Drosha* (mouse) | F | GAAGTCACCGTGGAGCTGAGTA |
| | R | ATCATTGCATGCTGACAGACATC |
| *Dicer* (mouse) | F | CAGCTCTGGACCATAACACAATTG |
| | R | AGGTCGCCCCTGATCTGAT |
| *Dgcr8* (mouse) | F | TCAAGGTCCGCCCTGTTTAT |
| | R | GAGGCACCAAAAGGCTCACTT |
| *Xpo5* (mouse) | F | GACGCAGAACATGGAAAGAATCT |
| | R | TGTCTTCATTTGTTGGTACTTGTTTACA |
| *Ago2* (mouse) | F | GCGTCAACAACATCCTGCT |
| | R | CTCCCAGGAAGATGACAGGT |
| *SRSF3* (human) | F | AACGGGCTTTTGGCTACTATG |
| | R | TTCACCATTCGACAGTTCCAC |
| *HPRT* (human) | F | CTGAGGATTTGGAAAGGGTGT |
| | R | GTAATCCAGCAGGTCAGCAAA |
| *CDKN1A*/p21 (human) | F | CAGCAGAGGAAGACCATGTG |
| | R | CGGCGTTTGGAGTGGTAGA |
| *TP53* (human) | F | AAGTCTGTGACTTGCACGTACTCC |
| | R | GTCATGTGCTGTGACTGCTTGTAG |
